# FORGEdb: a tool for identifying candidate functional variants and uncovering target genes and mechanisms for complex diseases

**DOI:** 10.1101/2022.11.14.516365

**Authors:** Charles E. Breeze, Eric Haugen, María Gutierrez-Arcelus, Xiaozheng Yao, Andrew Teschendorff, Stephan Beck, Ian Dunham, John Stamatoyannopoulos, Nora Franceschini, Mitchell J. Machiela, Sonja I. Berndt

**Affiliations:** Division of Cancer Epidemiology and Genetics, National Cancer Institute, National Institutes of Health, Bethesda, MD 20892; Altius Institute for Biomedical Sciences, 2211 Elliott Avenue 98121 Seattle; UCL Cancer Institute, University College London, 72 Huntley Street, London WC1E 6BT, United Kingdom; Division of Immunology, Department of Pediatrics, Boston Children’s Hospital, Harvard Medical School, Boston, MA, USA; Broad Institute of MIT and Harvard, Cambridge, MA, USA; CAS Key Lab of Computational Biology, CAS-MPG Partner Institute for Computational Biology, Shanghai Institute for Biological Sciences, Chinese Academy of Sciences, 320 Yue Yang Road, Shanghai 200031, China; European Molecular Biology Laboratory, European Bioinformatics Institute (EMBL-EBI), Wellcome Genome Campus, Hinxton, Cambridge, CB10 1SD, UK; Department of Epidemiology, University of North Carolina, Chapel Hill, NC, USA

## Abstract

The majority of disease-associated variants identified through genome-wide association studies (GWAS) are located outside of protein-coding regions and are overrepresented in sequences that regulate gene expression. Prioritizing candidate regulatory variants and potential biological mechanisms for further functional experiments, such as genome editing, can be challenging, especially in regions with a high number of variants in strong linkage disequilibrium or multiple proximal gene targets. Improved annotation of the regulatory genome can help identify promising variants and target genes for functional genomics experiments. To advance this area, we developed FORGEdb (https://forge2.altiusinstitute.org/files/forgedb.html), a web-based tool that can rapidly integrate data for individual genetic variants, providing information on associated regulatory elements, transcription factor (TF) binding sites and target genes for over 37 million variants. FORGEdb uses annotations derived from data across a wide range of biological samples to delineate the regulatory context for each variant at the cell type level. Multiple data types, such as Combined Annotation Dependent Depletion (CADD) scores, expression quantitative trait loci (eQTLs), activity-by-contact (ABC) interactions, Contextual Analysis of TF Occupancy (CATO) scores, transcription factor (TF) motifs, DNase I hotspots, histone mark ChIP-seq peaks and chromatin states, are included in FORGEdb and these annotations are integrated into a FORGEdb score to guide assessment of functional importance. In summary, FORGEdb provides an expansive and unique resource of genomic annotations and an integrated score that can be used to accelerate the translation of identified genetic loci into biological insight.

## Main

Genome-wide association studies (GWAS) have been remarkably successful in identifying genetic loci associated with many different diseases and traits [1]. As of the end of 2022, the GWAS catalog comprised >232,000 distinct variants associated with >3,000 diseases and traits [2]. Many loci identified from GWAS are intergenic or intronic, locating to non-protein-coding regions of the genome [3]. Although the functional mechanisms of some variants have been reported [4], most genomic loci have not been carefully studied and little is known about target genes, pathways or mechanisms of action. Multiple reports suggest that GWAS variants are overrepresented in sequences that regulate gene expression [3] [5]. Studies have shown enrichment for GWAS variants in cell- and tissue-specific regulatory elements [3] [5].

To aid interpretation of GWAS variants in the context of gene regulation, researchers have used large-scale mapping data for enhancers and other regulatory elements from ENCODE [6], Roadmap Epigenomics [7], and other consortia [8]. Several webtools, such as Haploreg [9], RegulomeDB [10] and others [11], have been developed to help researchers link these data to individual variants. However, these methods do not include more recent high-dimensional ENCODE data from contemporary technologies, such as Hi-C [12], or expanded expression quantitative trait locus (eQTL) data from large consortia, such as the Genotype-Tissue Expression Project (GTEx) [13] or the eQTLGen project [14]. Gathering relevant information from many different data sources and linking the data to individual genetic variants can be challenging in terms of computational resources, data processing, quality control, and reproducibility.

To address this issue and provide researchers with a state-of-the-art web tool for variant annotation that includes these updated resources, we developed FORGEdb (https://forge2.altiusinstitute.org/files/forgedb.html, **Table 1**). FORGEdb incorporates a range of datasets covering three broad areas relating to gene regulation: regulatory regions, transcription factor (TF) binding, and target genes. First, using genome-wide epigenomic track data from ENCODE [6], Roadmap Epigenomics [7], and BLUEPRINT [8] consortia, FORGEdb links SNPs with data for candidate regulatory regions (e.g., enhancers or promoters). Specifically, FORGEdb annotates variants for overlap with DNase I hotspots, histone mark broadPeaks, and chromatin states across a wide range of cell and tissue types. Second, within these candidate regulatory regions, FORGEdb integrates SNPs with transcription factor (TF) binding data via: a) the overlap with TF motifs and b) SNP-specific Contextual Analysis of TF Occupancy (CATO) scores, which provide a complementary line of evidence for TF binding computed from allele-specific TF occupancy data measured by DNase I footprinting[15]. Third, FORGEdb links SNPs to target genes by providing: a) the overlap between SNPs and enhancer-to-promoter looping regions (or other looping regions) using Activity-By-Contact (ABC) data [16] and b) expression quantitative trait locus (eQTL) annotations using large-scale data from GTEx [13] and eQTLGen [14]. In addition, FORGEdb includes annotations from datasets that aid interpretation of protein-coding changes. Specifically, it includes CADD scores, which measure the deleteriousness of SNPs using experimental data and simulated mutations [17]. By amalgamating these datasets into a single resource, FORGEdb offers an expanded set of annotations and a more comprehensive evaluation of individual variants beyond those provided by other commonly used webtools (**Table 1**) [9] [10] [11].

**Table 1:** A comparison of features across FORGEdb, Haploreg and RegulomeDB:

|  | FORGEdb | Haploreg | RegulomeDB |
| --- | --- | --- | --- |
| Roadmap Chromatin states | yes | yes | yes |
| TF motifs | yes | yes | yes |
| SNP Scoring system | yes | no | yes |
| Roadmap DNase-seq | yes | yes | no |
| Roadmap H3 histone mark data | yes | yes | no |
| 3D genomic data (ABC Hi-C-based data) | yes | no | no |
| CADD v1.6 data | yes | no | no |
| GTEx v8 data | yes | no | no |
| QTLGen data | yes | no | no |
| BLUEPRINT DNase-seq | yes | no | no |
| Allele-specific TF binding data (CATO) | yes | no | no |

To summarize the regulatory annotations and prioritize genetic variants for functional validation, we created a new scoring system for SNPs, combining all annotations relating to gene regulation into a single score called a FORGEdb score. In order to ensure that no single annotation or dataset would dominate or skew the scoring system, leading to bias, we adopted a points-based approach that evaluates each distinct experimental or technological approach separately. FORGEdb scores are computed based on the presence or absence of 5 independent lines of evidence for regulatory function:

1. DNase I hotspot, marking accessible chromatin (2 points)
2. Histone mark ChIP-seq broadPeak, denoting different regulatory states (2 points)
3. TF motif (1 point) and CATO score (1 point), marking potential TF binding
4. Activity-by-contact (ABC) interaction, indicating gene looping (2 points)
5. Expression quantitative trait locus (eQTL), demonstrating an association with gene expression (2 points)

FORGEdb scores are calculated by summing the number of points across all lines of evidence present for each SNP and range between 0 and 10. A score of 9 or 10 suggests a large amount of evidence for functional relevance, whereas 0 or 1 indicate a low amount of evidence. For example, there is evidence for eQTLs (for *IRX3* and *FTO*), chromatin looping, TF motifs, DNase I hotspots, and histone mark broadPeaks for rs1421085, a SNP previously identified for obesity[18] (**Figure 1**). Together, these annotations provide strong evidence for a regulatory role for this SNP with a FORGEdb score of 9. This high FORGEdb score for rs1421085 is consistent with independent experimental analyses that have demonstrated a regulatory role for this SNP with *IRX3* being a key target gene [4].

**Figure 1:**
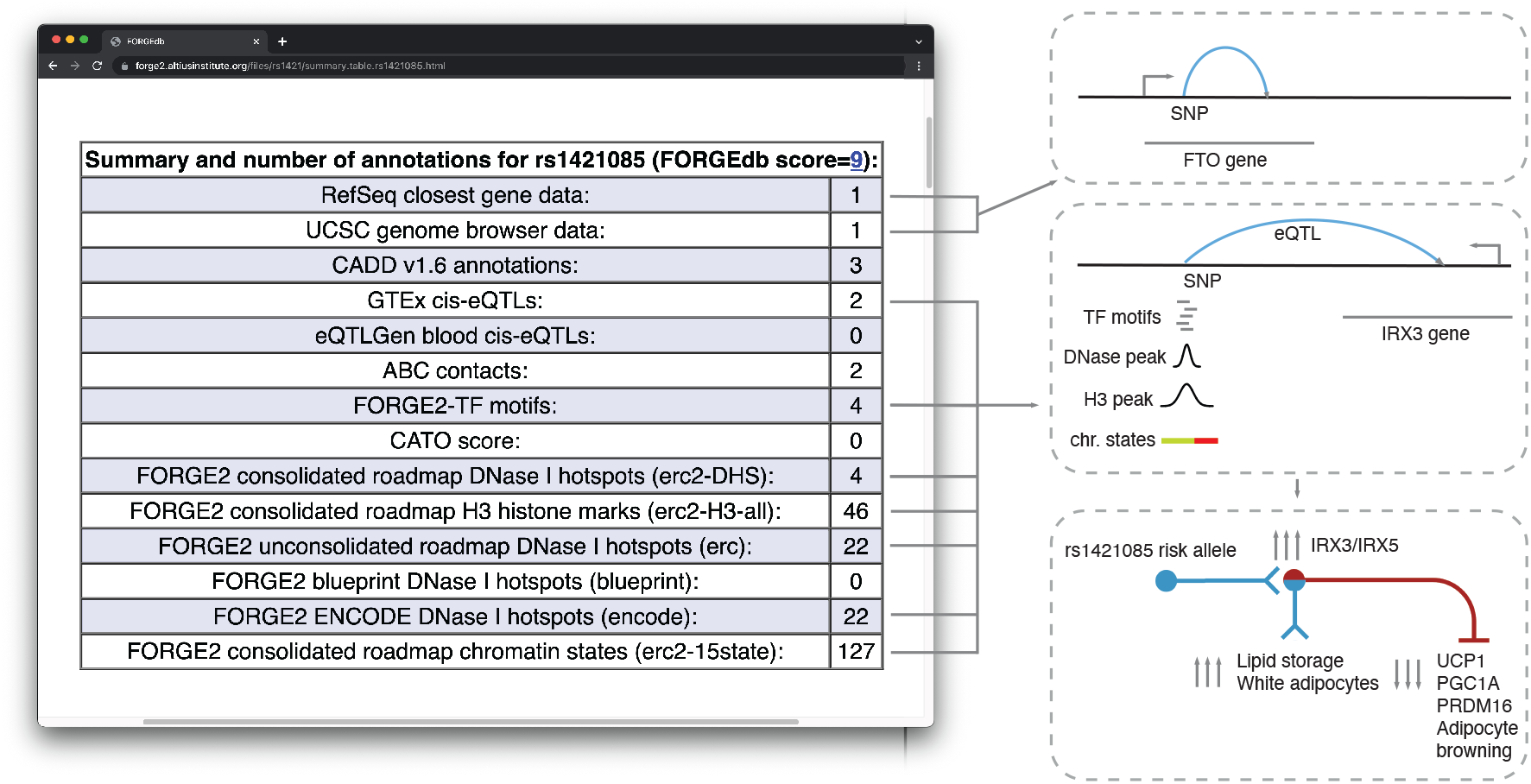
Example FORGEdb results for rs1421085. For this SNP, there is evidence for eQTL associations (with *IRX3* and *FTO*), chromatin looping (ABC interactions), overlap with significant TF motifs, DNase I hotspot overlap, as well as overlap with histone mark broadPeaks. The only regulatory dataset that this SNP does not have evidence for is for CATO score (1 point). The resulting FORGEdb score for rs1421085 is therefore 9 = 2 (eQTL) + 2 (ABC) + 1 (TF motif) + 2 (DNase I hotspot) + 2 (histone mark ChIP-seq). Independent experimental analyses by Claussnitzer *et al*. have demonstrated a regulatory role for this SNP in the regulation of white vs. beige adipocyte proliferation via *IRX3/IRX5* [4].

To assess the potential utility of FORGEdb scores in large-scale analysis of GWAS data, we obtained summary statistics from published studies of 30 traits/diseases (**Methods**) [2,19–41] and evaluated the correlation between FORGEdb scores and the ranking of SNPs by association p-value in each GWAS. Specifically, we binned the SNPs according to their association -log10 p-value and estimated the mean FORGEdb score for each bin. Results revealed a significant positive correlation between mean FORGEdb score and ranked SNP bins across all 30 phenotypes, with more significant p-values corresponding to higher FORGEdb scores (**Figure 2**, median correlation=0.845, range 0.55 to 0.98). As fine-mapping studies can identify sets of variants more likely to be functional, we compared FORGEdb scores for statistically-derived 95% credible set variants with FORGEdb scores for reported top SNPs from the same published study [42]. We discovered a significant overrepresentation of higher FORGEdb scores in the 95% credible sets (t-test p-value=0.002). These findings demonstrate that FORGEdb scores correlate with variants identified from GWAS and may be useful for prioritizing SNPs across a wide range of human traits and diseases, from common traits such as brown hair color and height to complex diseases like schizophrenia and lung cancer.

**Figure 2:**
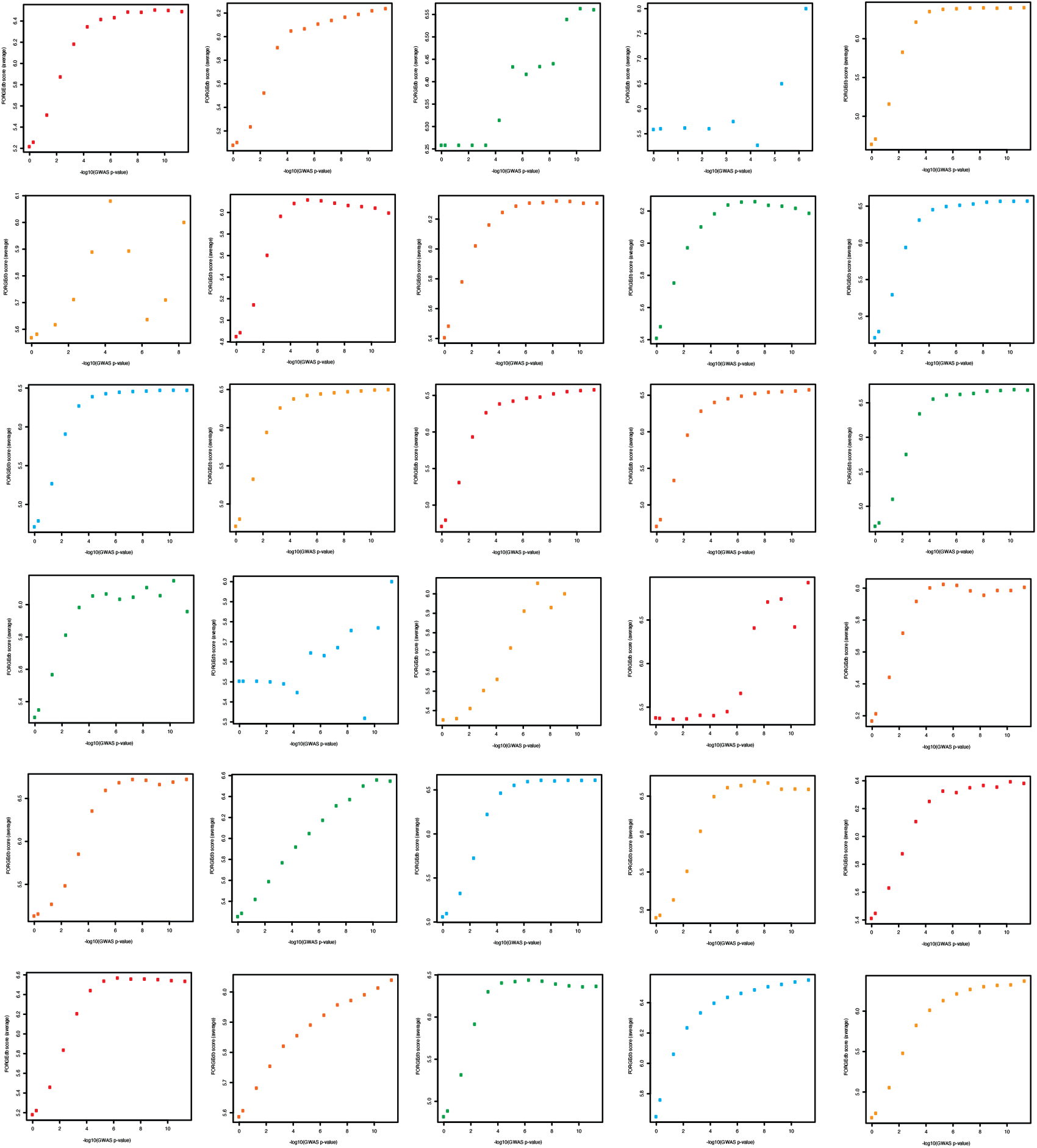
FORGEdb score (average, y-axis) versus GWAS -log10(p-value) (x-axis) across 30 GWAS. Each point shows the FORGEdb score average across all GWAS SNPs at a each p-value cutoff. From top left to bottom right: venous thrombosis (cor=0.87), hair colour (cor=0.89), melanoma (cor=0.95), colon cancer (cor=0.69), LDL (cor=0.81), fasting insulin (cor=0.55), prostate cancer (cor=0.78), diastolic blood pressure (cor=0.82), systolic blood pressure (cor=0.80), neutrophil cell count (cor=0.82), eosinophil cell count (cor=0.81), lymphocyte cell count (cor=0.82), monocyte cell count (cor=0.84), white blood cell count (cor=0.83), basophil cell count (cor=0.85), estimated glomerular filtration rate (cor=0.79), major depressive disorder (cor=0.59), autism (cor=0.96), attention deficit hyperactivity disorder (cor=0.89), breast cancer (cor=0.79), lung cancer (cor=0.90), schizophrenia (cor=0.98), rheumatoid arthritis (cor=0.86), Alzheimer’s disease (cor=0.86), type 2 diabetes (cor=0.88), inflammatory bowel disease (cor=0.85), body mass index (cor=0.96), red blood cell count (cor=0.77), height (cor=0.86), and waist to hip ratio (cor=0.90).

To assess the utility of FORGEdb scores in identifying potential functional variants, we examined the relationship between FORGEdb scores and expression-modulating variants (emVars), which are candidate functional variants prioritized from massively parallel reporter assays (MPRAs). We utilized emVar data from Tewhey *et al*.[43], who evaluated 39,487 variants using MPRAs, identifying 248 variants that had a high effect on gene expression. Comparing these 248 emVars with 37 million variants in the FORGEdb database revealed a significant overrepresentation of emVars in higher FORGEdb scores (paired t-test p-value=0.005, **Figure 3**). This indicates that variants with higher FORGEdb scores are more likely to be functional and that FORGEdb scores are likely well-suited for prioritizing variants in MPRAs and other massively parallel experiments. Moreover, emVars exhibited significantly higher FORGEdb scores than 39,487 candidate MPRA variants from the same study (paired t-test p-value=0.004), suggesting that FORGEdb scores add further information not present in previous variant prioritization methods.

**Figure 3:**
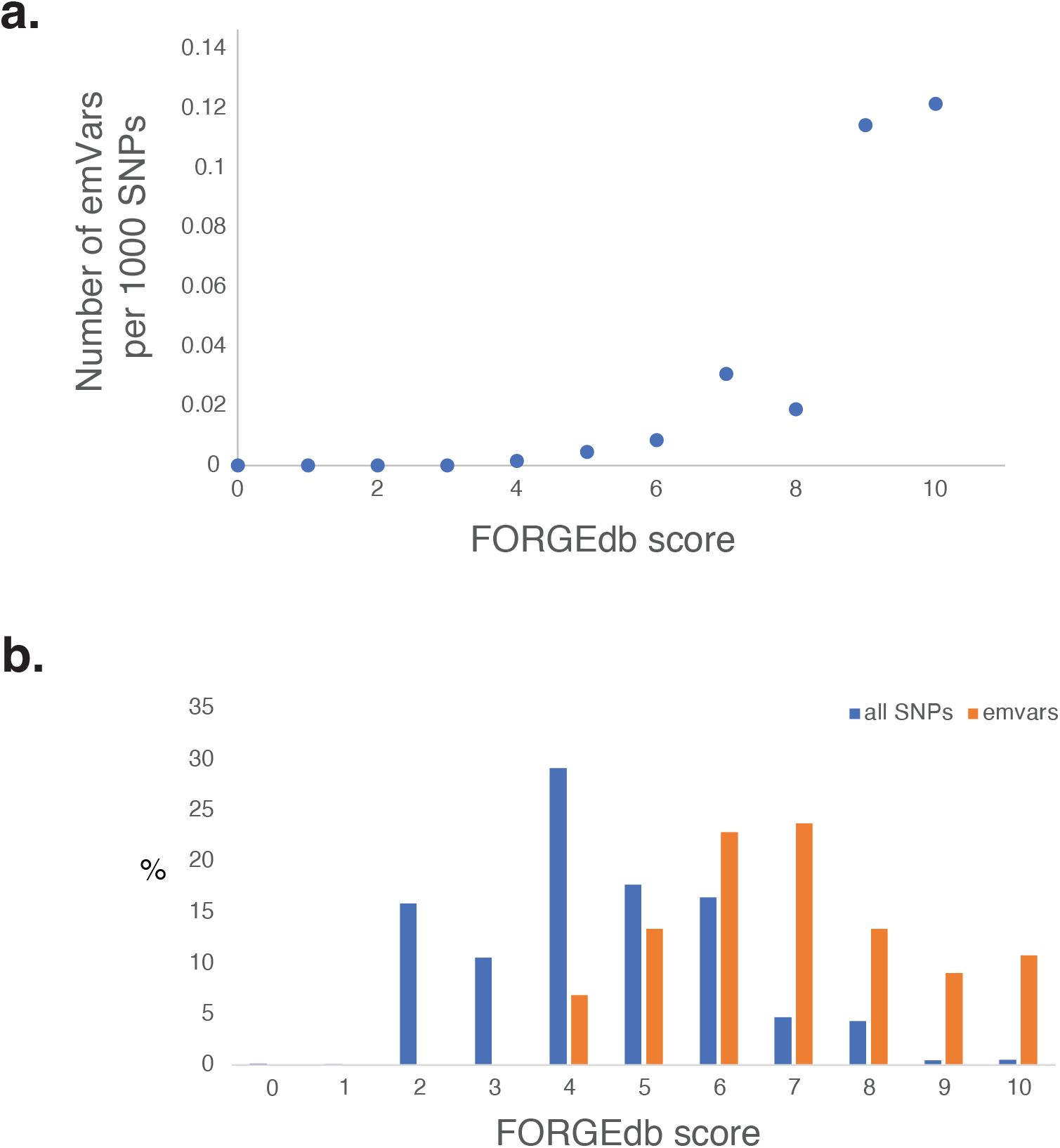
Expression-modulating variants (emVars) identified from massively parallel reporter assays (MPRAs)[43] are overrepresented in top FORGEdb scores. Shown here are: a. the number of emVars per 1000 SNPs with the same FORGEdb score (y-axis) for each possible FORGEdb score (0-10) (x-axis) and b. a histogram of FORGEdb scores for emVars (orange) and ∼37 million SNPs available in FORGEdb (blue).

In summary, FORGEdb is a new web-based tool to aid the interpretation and prioritization of genetic variants for experimental analysis. FORGEdb includes a number of features from novel technologies not available in commonly used webtools, providing a more comprehensive analysis of potential regulatory function [9] [10] [11]. All of these features are accessible via a simple, easy-to-use search engine that can be found at https://forge2.altiusinstitute.org/files/forgedb.html. Annotations from FORGEdb can be accessed from https://ldlink.nci.nih.gov/?tab=ldproxy, https://ldlink.nci.nih.gov/?tab=ldassoc, https://ldlink.nci.nih.gov/?tab=ldmatrix, and https://forge2.altiusinstitute.org/ [5] [44] [45].

## Methods

### Databases used in FORGEdb

FORGEdb standalone first annotates variants for overlap with DNase I hotspots, histone mark broadPeaks, and chromatin states across a wide range of cell and tissue types as implemented in FORGE2 [5]. Second, FORGEdb annotates variants for Activity-By-Contact (ABC) data (as implemented in Fulco *et al*.) [16], Contextual Analysis of TF Occupancy (CATO) scores as implemented by Maurano *et al*. [15], TF motifs as implemented in FORGE2-TF (https://forge2-tf.altiusinstitute.org/), significant eQTLs from GTEx [13] and eQTLGen [14], and closest gene from RefSeq[46].

### Accessing FORGEdb

FORGEdb is available through a web browser (https://forge2.altiusinstitute.org/files/forgedb.html). A programmatic inteface to FORGEdb has been developed via CRAN package LDlinkR (https://cran.r-project.org/web/packages/LDlinkR/index.html). FORGEdb code is available from https://github.com/charlesbreeze/FORGEdb. FORGEdb databases can be downloaded from https://forgedb.altiusinstitute.org/?download.

### Example analysis

Briefly, FORGEdb uses a Common Gateway Interface (CGI) perl-based web server, which returns HTML pages from search queries at https://forge2.altiusinstitute.org/files/forgedb.html. An example FORGEdb analysis is available at https://forge2.altiusinstitute.org/files/rs1421/summary.table.rs1421085.html and a description of FORGEdb scores is available at https://forge2.altiusinstitute.org/files/FORGEdb_scores.html. A brief tutorial on FORGEdb is avilable at https://github.com/charlesbreeze/FORGEdb#readme.

### Regenerating FORGEdb pages

To regenerate FORGEdb HTML page component using local SSD memory on a distributed high-performance computing system we also provide guidelines and code at https://github.com/charlesbreeze/FORGEdb. Note that computational requirements will be relatively large and use of local memory is preferred, a file accesses can run into the many millions of database queries per second.

### Analysis of MPRA and GWAS data

For analysis of MPRA emVar data, we downloaded the SNP information from table S1 (39,478 ref/alt pairs tested by MPRA) and S2 (emVars) of Tewhey *et al*., 2016 [43]. We then generated FORGEdb SNP scores for all 248 reported emVars and the other 39,478 SNPs evaluated in the manuscript. We then compared the FORGEdb scores for the emVars with the other evaluated SNPs and 37 million SNPs available in FORGEdb.

For analysis of GWAS data across 30 disease/traits, we downloaded summary statistics from OpenGWAS [20] and other sources [2,19–41]. Ethnicities analysed in these GWAS include African American/Afro-Caribbean, East Asian and European. For each GWAS, we computed FORGEdb scores across all variants and then computed the average score at different p-value thresholds. Published 95% credible sets for a coronary heart disease GWAS were obtained from van der Harst et al. [42]. Plotting and statistical analyses were conducted in R [47].

### Contact

For any questions or information on HTML component generation contact.

## Acknowledgements

This study was supported in part by the Intramural Research Program of the Division of Cancer Epidemiology and Genetics, National Cancer Institute, NIH. We would like to acknowledge the IHEC Integrative analysis project for supporting this research. SB acknowledges funding from the Wellcome Trust (218274/Z/19/Z).

## Availability of data and materials

FORGEdb code is available from https://github.com/charlesbreeze/FORGEdb. FORGEdb databases can be downloaded from https://forgedb.altiusinstitute.org/?download.

## Appendix

### Tutorial: generating 37M FORGEdb pages

This tutorial lists the different steps to generate 37 million webpages using FORGEdb standalone. The code presented here is optimised for the NIH Biowulf cluster, but can be amended to work on other large compute clusters. Note that different required FORGEdb databases can be downloaded from https://forge2.altiusinstitute.org/?download.

### Tutorial part 1: Preparatory steps

The “file.commands.regenerate.forgedb.htmls.sh.using.local.memory.sh” is a file containing all forgedb standalone commands for each SNP, one per line. There are 37964763 SNPs

~~~
       [breezece@biowulf breezece]$ wc
file.commands.regenerate.forgedb.htmls.sh.using.local.memory.sh &
       [3] 68788
       [breezece@biowulf breezece]$ 37964763
file.commands.regenerate.forgedb.htmls.sh.using.local.memory.sh
~~~

To split into 312 nodes:

~~~
       [breezece@biowulf breezece]$ bc
       37964763/312
       121681.93269230769230769230
       quit
~~~

We then split the file:

~~~
        [breezece@biowulf breezece]$ split -121682
file.commands.regenerate.forgedb.htmls.sh.using.local.memory.sh
test.localmem2/for.job &
        [3] 69391
        breezece@biowulf breezece]$
        [3]+ Done    split-121682
file.commands.regenerate.forgedb.htmls.sh.using.local.memory.sh
test.localmem2/for.job
~~~

List all the files made:

~~~
        [breezece@biowulf breezece]$ ls test.localmem2/ > ls.localmem
        [breezece@biowulf breezece]$ wc ls.localmem
        312 ls.localmem
~~~

Modify myjobscript template to include the real files from each directory:

~~~
      [breezece@biowulf breezece]$ cat ls.localmem |awk ‘{print “cat myjobscript|sed
\”s/test.sh/”$0”/g\” > test.localmem2/myjobscript.”$0}’|bash
~~~

### Tutorial part 2: running code on cluster

Run with (from directory /data/breezece/test.localmem2):

~~~
sbatch --cpus-per-task=32 --mem=120g --gres=lscratch:700 --constraint=x2650 --partition norm --time=9-03:08:42 myjobscript.for.jobaa
~~~

(to run all at once the commands are):

~~~
ls myjob* |grep -v head|awk ‘{print “sbatch --cpus-per-task=32 --mem=120g --gres=lscratch:700 --constraint=x2650 --partition norm --time=9-03:08:42 “$0}’ > submits.jobs.sbatch.sh
[test.localmem2]$ nohup bash submits.jobs.sbatch.sh &
~~~

this runs the modified version of myjobscript:

~~~
      [breezece@biowulf breezece]$ cat test.localmem7/myjobscript.for.jobaa
      #!/bin/bash
      mkdir /lscratch/$SLURM_JOBID/done.htmls
      cp for.jobaa /lscratch/$SLURM_JOBID
      cd /lscratch/$SLURM_JOBID
      cp /data/breezece/tableToHtml2.pl /lscratch/$SLURM_JOBID
      cp /data/breezece/miniabout.txt /lscratch/$SLURM_JOBID
      cp /data/breezece/img.txt /lscratch/$SLURM_JOBID
      cp /data/breezece/tableToHtml.pl /lscratch/$SLURM_JOBID
      cp /data/breezece/minibr.txt /lscratch/$SLURM_JOBID
      cp /data/breezece/get.table.snp.forge2.2.sh /lscratch/$SLURM_JOBID
      cp /data/breezece/abc.db /lscratch/$SLURM_JOBID
      
      cp /data/breezece/cato.db /lscratch/$SLURM_JOBID
      cp /data/breezece/closest.refseq.gene.hg19.db /lscratch/$SLURM_JOBID
      cp /data/breezece/eqtlgen.db /lscratch/$SLURM_JOBID
      cp /data/breezece/forge_2.0.db /lscratch/$SLURM_JOBID
      cp /data/breezece/forge2.tf.fimo.jaspar.1e-5.taipale.1e-5.taipaledimer.1e-5.uniprobe.1e-5.xfac.1e-5.db /lscratch/$SLURM_JOBID
      cp /data/breezece/gtex.eqtl.db /lscratch/$SLURM_JOBID
      cp /data/breezece/dmp_GWAS_bins /lscratch/$SLURM_JOBID
      cp /data/breezece/dmp_GWAS_params /lscratch/$SLURM_JOBID

      cp /data/breezece/snps.to.avoid /lscratch/$SLURM_JOBID
      cp /data/breezece/scores.py /lscratch/$SLURM_JOBID
      
      export SLURM_JOBID
      # for.jobaa should run, say, 100 commands
      #cat for.jobaa |/usr/local/apps/parallel/20220422/bin/parallel -j
$SLURM_CPUS_PER_TASK
      cat for.jobaa |/usr/local/apps/parallel/20220422/bin/parallel -j 16

      # after all the parallel jobs have finished
      
      tar -cvz -f $SLURM_JOBID.done.htmls.tar.gz
/lscratch/$SLURM_JOBID/done.htmls

      cp $SLURM_JOBID.done.htmls.tar.gz /data/breezece/done.htmls
and for.jobaa
      [breezece@biowulf breezece]$ head test.localmem7/for.jobaa
      TMPDIR=/lscratch/$SLURM_JOBID; bash get.table.snp.forge2.2.sh rs10000242
      TMPDIR=/lscratch/$SLURM_JOBID; bash get.table.snp.forge2.2.sh rs10000384
~~~

After jobs are complete, the full list of 37M webpages (in compressed format) can be viewed at the “done.htmls” directory.

